# Structure of the *Staphylococcus aureus* bacteriophage 80α neck shows the interactions between DNA, tail completion protein and tape measure protein

**DOI:** 10.1101/2024.12.10.627806

**Authors:** James L. Kizziah, Amarshi Mukherjee, Laura K. Parker, Terje Dokland

**Affiliations:** Department of Microbiology, University of Alabama at Birmingham, Birmingham, AL 35294, USA

**Keywords:** cryo-electron microscopy, 3D reconstruction, virus capsid, *Staphylococcus aureus* pathogenicity island 1, bacteriophage tail, portal protein, connector

## Abstract

Tailed bacteriophages with double-stranded DNA genomes (class *Caudoviricetes*) play an important role in the evolution of bacterial pathogenicity, both as carriers of genes encoding virulence factors and as the main means of horizontal transfer of mobile genetic elements (MGEs) in many bacteria, such as *Staphylococcus aureus*. The *S. aureus* pathogenicity islands (SaPIs), including SaPI1, are a type of MGEs are that carry a variable complement of genes encoding virulence factors. SaPI1 is mobilized at high frequency by “helper” bacteriophages, such as 80α, leading to packaging of the SaPI1 genome into virions made from structural proteins supplied by the helper. 80α and SaPI1 virions consist of an icosahedral head (capsid) connected via a unique vertex to a long, non-contractile tail. At one end of the tail, proteins associated with the baseplate recognize and bind to the host. At the other end, a connector or “neck” forms the interface between the tail and the head. The neck consists of several specialized proteins with specific roles in DNA packaging, phage assembly, and DNA ejection. Using cryo-electron microscopy and three-dimensional reconstruction, we have determined the high-resolution structure of the neck section of SaPI1 virions made in the presence of phage 80α, including the dodecameric portal (80α gene product (gp) 42) and head-tail-connector (gp49) proteins, the hexameric head-tail joining (gp50) and tail terminator (gp52) proteins, and the major tail protein (gp53) itself. We were also able to resolve the DNA, the tail completion protein (gp51) and the tape measure protein (gp56) inside the tail. This is the first detailed structural description of these features in a bacteriophage, providing insights into the assembly and infection process in this important group of MGEs and their helper bacteriophages.

## INTRODUCTION

Tailed bacteriophages (phages)—viruses that infect bacteria—with double-stranded DNA genomes belonging to class *Caudoviricetes* (order *Caudovirales*) are abundant in all environments and play important roles in biomass turnover and bacterial evolution [1]. Phages often exist as prophages, integrated as mobile genetic elements (MGEs) into their host genomes, and frequently carry genes encoding virulence factors and antibiotic resistance [2-3].

The *Caudovirales* phages consist of an icosahedral or prolate head (capsid) attached to a tail via a unique vertex that incorporates a dodecameric portal. The portal is assumed to act as the nucleus for capsid assembly, and serves as an entry and exit point for the DNA [4–6]. Bacteriophages are traditionally divided into groups based on tail morphology: siphoviruses have long, non-contractile tails, myoviruses have long contractile tails, while podoviruses have short tails. However, these groups are not monophyletic and are no longer considered viral families [7]. The tail generally makes the first contact with the host and play important roles in host recognition, breakdown of the cell wall, and injection of the genome [8–10]. Long tails are assembled through a separate pathway and are attached to heads via a connector or neck region, consisting of several proteins that together act as a conduit for the DNA during infection [11].

*Staphylococcus aureus* is an opportunistic human pathogen that encodes a large array of virulence factors [12,13]. Most of these are encoded on MGEs, including prophages and chromosomal islands [14,15]. Transduction by phages represents the main mechanism by which MGEs are transferred between hosts in *S. aureus* [16,17]. *S. aureus* pathogenicity islands (SaPIs) are a type MGEs that encode superantigen toxins, adhesins and other virulence factors. SaPIs become mobilized at high frequency by so-called “helper” phages [18–20] and packaged into transducing particles made up of phage-encoded structural proteins. SaPIs suppress the replication of their helpers, often by redirecting the assembly pathway to form capsids of a smaller size than that normally made by the phage [20,21].

Phage 80α is a typical temperate staphylococcal siphovirus that can act as helper for a number of SaPIs, including SaPI1 [19]. Phages closely related to 80α are found in a wide variety of staphylococcal strains, including MRSA strain USA300 LAC, commonly involved in community-acquired infections [22]. The 80α virion has a 63 nm icosahedral head with *T*=7 architecture, a 190 nm long flexuous tail, a complex baseplate [23–25], and a 43,864 base pair double-stranded (ds)DNA genome [26](NCBI RefSeq NC_009526.1)(Fig. 1A). When 80α acts as a helper for SaPI1, the assembly pathway is redirected to form small capsids with *T*=4 architecture [27,28] (Fig. 1B). Both 80α and SaPI1 genomes are packaged via a headful mechanism yielding blunt-ended, circularly permuted copies with terminal sequences distributed throughout the genome [29].

**Figure 1.**
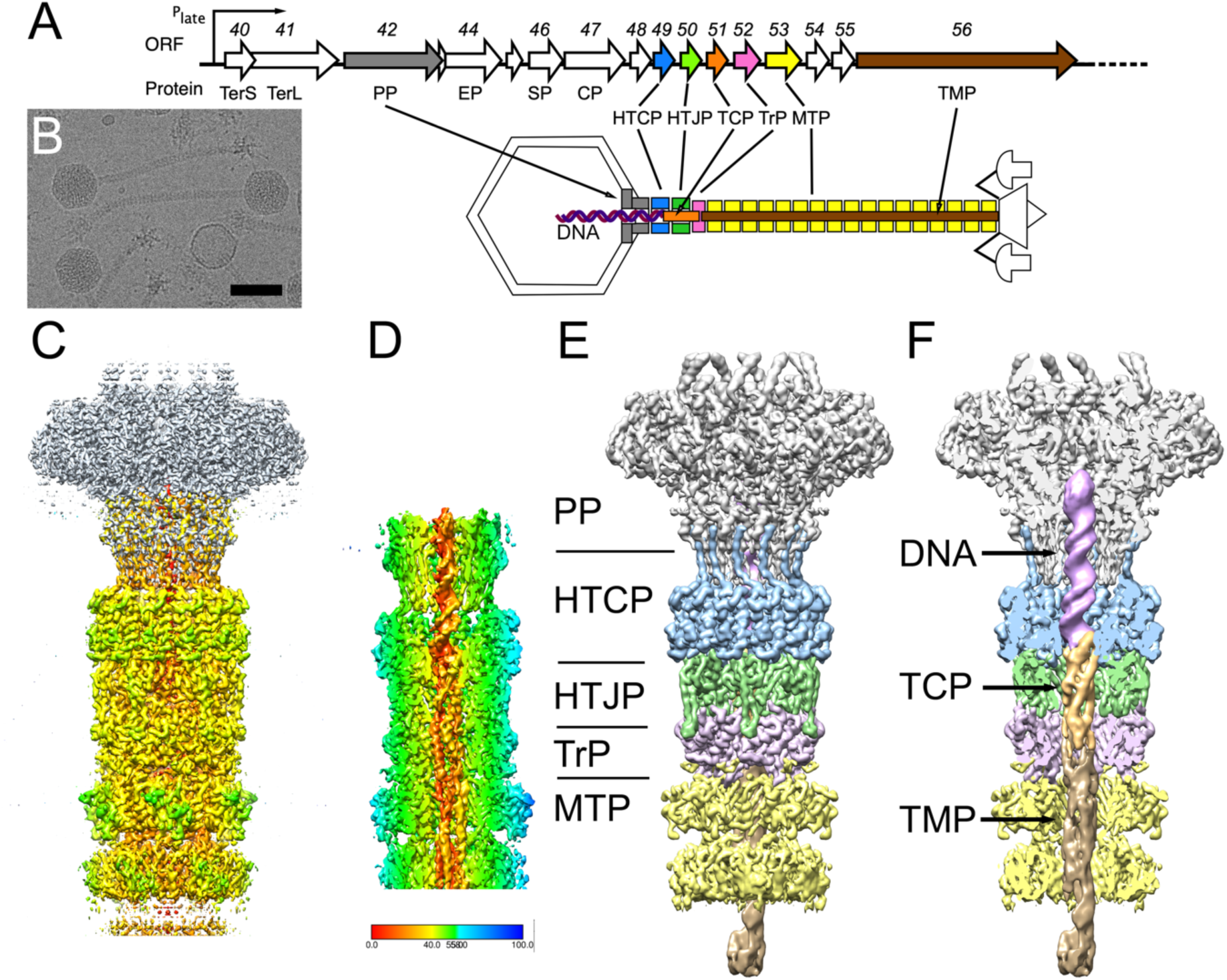
Structure of the SaPI1 neck. (A) Schematic diagram of part of the late operon of 80α, showing the genes encoding neck proteins. ORF numbers and protein names are shown: TerS, small terminase subunit; TerL, large terminase subunit; PP, portal protein; EP, ejection protein, SP, scaffolding protein; CP, major capsid protein; HTCP, head-tail connector protein; HTJP, head-tail joining protein; TCP, tail completion protein; TrP, tail terminator protein; MTP, major tail protein; TMP, tape measure protein. The schematic diagram indicates the location of the neck proteins in the virion. The DNA is shown as a purple double helix. (B) Cryo-electron micrograph of SaPI1 virions. Scale bar = 50 nm. (C) Isosurface representation of a composite of the C6 neck reconstruction, colored by radius from the central axis, and the previously determined C12 reconstruction of the portal (gray) [30]. (D) Cutaway view of the asymmetric (C1) reconstruction of the neck, showing the DNA, TCP and TMP inside the neck, colored by radius according to the color bar. (E) Segmented reconstruction, colored by protein as in panel A: PP (gp42), gray; HTCP (gp49), blue; HTJP (gp50), green; TrP (gp52), pink; MTP (gp53), yellow. (F) Cutaway view of segmented reconstruction, showing density inside neck, colored as in panel C. In addition, the DNA is purple; TCP (gp51), orange; TMP (gp56), brown.

We previously determined structures of 80α and SaPI1 procapsids and mature capsids [24,28], as well as the 80α portal protein expressed in *E. coli* and in situ in SaPI1 virions [30]. We also determined the structure of the 80α baseplate, a complex of 7 different proteins, including three presumed receptor binding proteins [25]. However, the organization of other tail-related components, including the neck that connects between the tail and the head was not resolved in these structures. Here, we have used cryo-electron microscopy (cryo-EM) to determine the structure of the neck region of SaPI1 virions that constitutes the interface between the head and the tail. The reconstruction resolves several ring-like proteins that connect the portal to the major tail protein. We were also able to resolve in molecular detail the DNA, the tape measure protein and the so-called “tail completion protein” inside the tail. This is the first detailed structural description of these features in a siphovirus, providing novel insights into the architecture and assembly of phage tails, and a structural basis for understanding the infection process in this important group of bacteriophages.

## RESULTS

### 1. The head-to-tail connector

Immediately below the portal there is a dodecameric ring that we previously identified as made from gp49, product of open reading frame (ORF) 49 in the RefSeq entry for 80α (NC_009526.1) (Fig. 1A). We refer to this as the “head-to-tail connector protein” (HTCP), consistent with usage in phage lambda (Table 1). In *Bacillus subtilis* phage SPP1 and *Pseudomonas aeruginosa* phage JBD30 it is referred to as an “adaptor”, as it connects between the dodecameric portal and the hexameric proteins below it [31,32].

**Table 1.** List of 80α neck proteins and equivalent proteins from other phages.

| 80α (*S. aureus*) | | | | Lambda (*E. coli*) [43,45]<br>[57] | | SPP1 (*B. subtilis*)<br>[31,39,42] | | HK97 (*E. coli*) [34,58,59] | | JBD30 (*P. aeruginosa*) [32] | | GTA [38]. | | Casjens<br>review [35] |
| --- | --- | --- | --- | --- | --- | --- | --- | --- | --- | --- | --- | --- | --- | --- |
| Protein | Abbr<sup>¶</sup> | Olig<sup>§</sup> | ORF | Name | ORF | Name | ORF | Name | ORF | Name | ORF | Name | ORF | Abbr |
| **HEAD ASSOCIATED:** |  |  |  |  |  |  |  |  |  |  |  |  |  |  |
| Portal | PP | 12 | 42 | Portal | B | Portal | 6 | Portal | 3 | Portal | 32 | Portal | Rcc01684 | – |
| Head-Tail<br>Connector | HTCP | 12 | 49 | Connector | W | Adaptor | 15 | Adaptor, Ad1;<br>Connector | 6 | Adaptor | 41 | Adaptor | Rcc01688 | HTJW/HTJ1 |
| Head-Tail Joining | HTJP | 6 | 50 | Head-tail joining | FII | Stopper | 16 | Head closure, Hc1;<br>Adaptor | 7 | Stopper* | 42* | Stopper | Rcc01689 | HTJ2 |
| **TAIL ASSOCIATED:** |  |  |  |  |  |  |  |  |  |  |  |  |  |  |
| Tail Completion | TCP | 1 | 51 | Tail completion | Z | TCP | 16.1 | – | 10 | Stopper* | 42* | – | – | TCP |
| Tail Terminator | TrP | 6 | 52 | Tail terminator | U | THJP | 17 | Tail terminator | 11 | – | – | Terminator | Rcc01690 | TrP |
| Major Tail | MTP | 6 | 53 | Tail tube | V | TTP | 17.1 | MTP | 12 | MTP | 44 |  |  | TTP |
| Tape measure | TMP | 3 | 55 | TMP | H | TMP | 18 | TMP | 16 |  |  |  |  | TMP |

¶Additional abbreviations not defined in this column: THJP, tail-head joining protein; TTP, tail tube protein; HTJW, head-tail joining protein, lambda gpW type.
§Oligomeric state of protein
\*Identified as “stopper” in Ref. 32 [32], but is actually equivalent to TrP.

In the focused reconstruction of the in situ portal [30], the C-terminus of the HTCP could be seen to add a β-strand to the clip domain of PP, forming a three-stranded β-sheet together with one strand from each of two adjacent PP subunits. However, the rest of the HTCP was not well resolved in the previous structure, and the density had been truncated due to the applied mask. We therefore re-extracted the particles from the same data set (Fig. 1B), centered on the neck region below the portal, and carried out a separate focused reconstruction that included part of the portal, the HTCP and additional structures associated with the neck and tail. While the HTCP was expected to have C12 symmetry, the tail is known to have C6 symmetry [25]; therefore, the focused reconstruction was refined with the application of C6 symmetry, reaching a final resolution of 3.1 Å (Fig. 1C; Fig. S1; Table S1).

A model for gp49 was generated using AlphaFold2 [33] and fitted into the map density immediately below the portal, followed by refinement in ISOLDE (Fig. 1E, 2A-E). The HTCP forms a dodecamer in which each subunit consists of a short N-terminal α-helix followed by three 25 Å-long antiparallel α-helices interspersed with a β-hairpin (Fig. 2A, F). The previously described extended C-terminal β-strand that inserts into the clip domain of PP matched well when an overlapping region consisting of the top of the last α-helix from the two models were merged (Fig. 2A-C, F). The three long α-helices from all 12 subunits form a 95 Å-wide outer ring, while the β-hairpins define a barrel with 31 Å inner diameter that lines the central tube of the neck, constituting the narrowest point of the neck below the portal (Fig. 2D, E). (The portal itself has an inner diameter of 26 Å at its narrowest [30].) The β-barrel inserts into the tube formed by the other neck proteins below (Fig. 2D, E).

**Figure 2.**
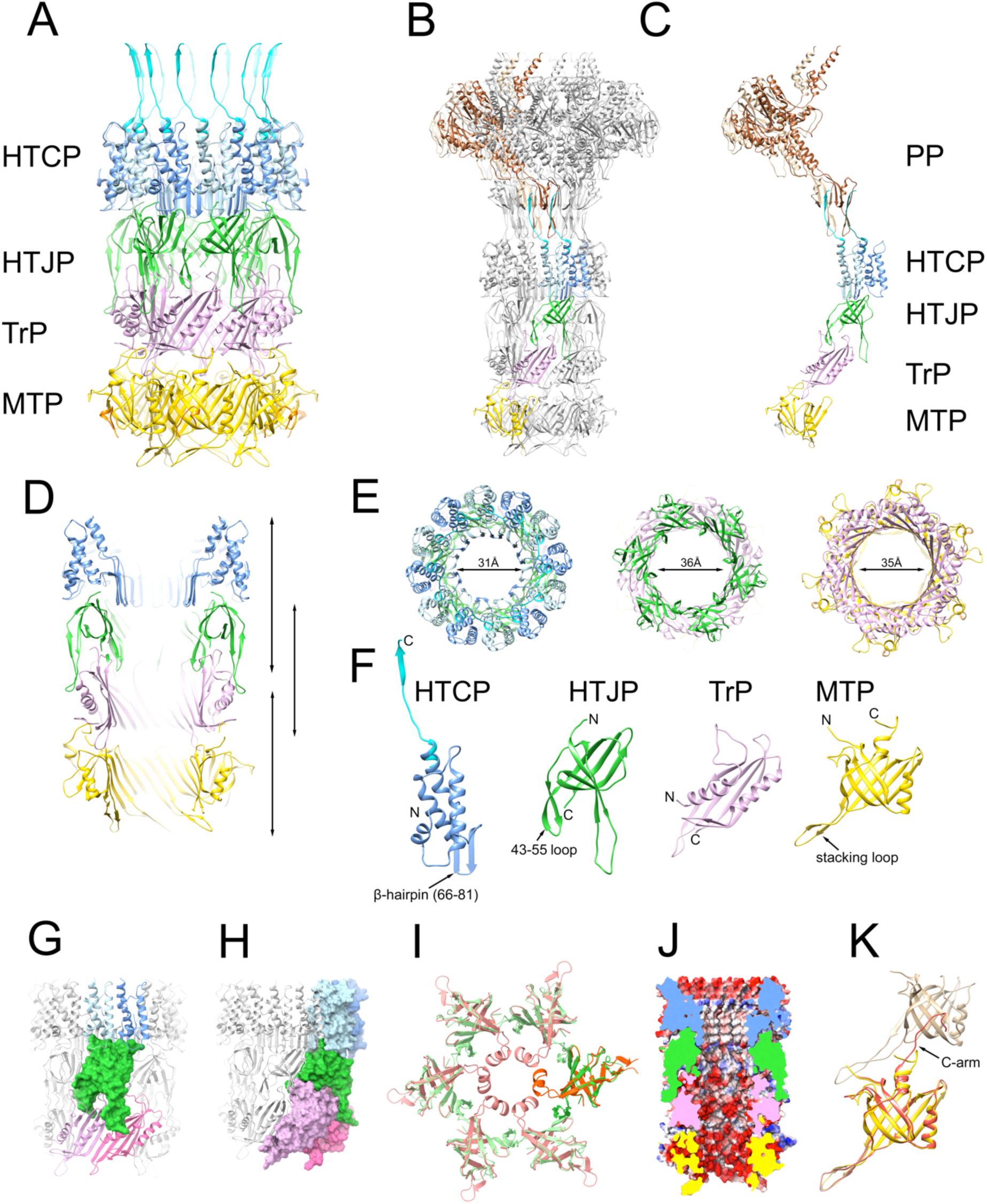
Atomic modeling of the neck. (A) Ribbon representation of the atomic model from the C6 neck reconstruction. The two copies of HTCP (gp49) per asymmetric unit are colored light and dark blue; the HTJP (gp50), TrP (gp52) and MTP (gp53) hexamers are shown in green, pink and yellow, respectively. The C-terminus of HTCP, which was built from the previous portal structure [30] is in cyan. (B) Atomic model of the neck reconstruction with one asymmetric unit colored as in panel A, the rest gray. The previously determined portal protein model is included, with the two copies of PP (gp42) per asymmetric unit colored light and dark brown. (C) An isolated asymmetric unit, colored as in A and B. (D) Slab through the neck model, colored as in panel A. (E) Sections through the neck model at the levels indicated by the arrows in panel D. From left to right: HTCP and HTJP; HTJP and TrP; TrP and MTP. The smallest inner diameters are indicated. (F) Monomers of neck proteins, from left to right: HTCP, HTJP, TrP and MTP. (G) Interaction between HTJP (green surface) and the HTCP (blue) and TrP (pink). (H) Interaction between HTJP (green), HTCP (blue) and TrP (pink) shown as molecular surfaces. (I) Cutaway view showing the electrostatic potential surface of the tail interior. (J) AlphaFold model of HTJP (orange) superimposed on the HTJP model from the neck reconstruction (green), showing the predicted inner ring of α-helices. (K) MTPs from the 80α baseplate reconstruction (orange and tan) superimposed on the MTP from the neck model (yellow), showing the difference in the C-arm.

The structure of gp49 is similar to the equivalent proteins from other phages, including gp6 from *Escherichia coli* phage HK97 [34], gp15 from phage SPP1 [31] and gp41 from JBD30, to which it aligns with root-mean-square deviations (RMSD) of 3.61Å, 4.84Å and 5.21Å, respectively (Fig. S2; Table S2). By comparison, the HTCP from *E. coli* phage lambda, gpW, is shorter, lacking the N-terminal two α-helices (Fig. S2). The remaining two α-helices of gpW are tilted ≈45° outward compared to 80α. However, the β-hairpin that lines the interior of the 80α and SPP1 HTCPs, but is missing in HK97 and JBD30, is retained in lambda, indicating that the proteins are indeed structurally and evolutionarily related [35]. In the solution structure of gpW [36], the hairpin is folded over the helices, suggesting that the protein undergoes a conformational change upon oligomerization and portal binding.

### 2. The head-to-tail joining protein

The focused reconstruction of the neck revealed two additional protein rings with C6 symmetry between the HTCP and the major tail protein (MTP, gp53) (Fig. 1C, E). Based on the principle of co-linearity of genome and structure observed in other phages, we presumed that these proteins corresponded to gp50, gp51, and/or gp52 (Fig. 1A), and made AlphaFold2 models for each protein. Each model was then tested for fit into the ring-like densities. While gp51 did not match either ring, gp50 and gp52 matched the top and bottom ring, respectively (Fig. 1E, 2A-E).

The gp50 protein, which makes up the topmost ring, is equivalent to the head-tail joining protein (HTJP) gpFII of lambda [37] and gp16 of SPP1 [31], as well as a protein (Rcc01689) from a phage-like “gene transfer agent” (GTA) from *Rhodobacter capsulatus* [38] (Fig. 2F; Fig. S2; Table S2). In SPP1, gp16 was associated with heads extracted from tail-less DNA-filled capsids [31] and was considered part of the head-tail connector complex, assumed to be added to packaged heads prior to attachment of the tail. SPP1 gp16 was designated as “stopper”, but whether it serves a functional role as a stopper—to prevent the DNA from falling out of the capsids after packaging—in 80α is not known. We will refer to this protein by the term “head-tail-joining protein” (HTJP), consistent with the terminology used by Casjens et al. [35] (Table 1).

The 80α HTJP consists of a seven-stranded β-sandwich with two extended loops (Fig. 2F). The top of the sandwich inserts into a groove between the outer α-helix and the β-hairpins formed by two copies of HTCP (Fig. 2D,G,H). The two extended loops at the other side of the sandwich form a clamp that grabs the inside and outside of the ring below it, made of gp52 (Fig. 2D,G,H; see below). Though the loop comprised of residues 43-55 exhibits a β-hairpin structure in the reconstruction, AlphaFold predicted residues 43-50 to be an α-helix that would narrow the inner diameter of the hexamer to 16 Å (Fig. 2I), possibly representing a pre-tail joining state where the protein ring could act as a stopper, similar to SPP1 [31].

The β-sandwich of HTJP resembles a classic “tail tube protein” (TTP) fold that is found in the major tail protein (MTP, gp53), as well as the distal tail protein (Dit) and the tail tip protein (Tal) [25]. The β-hairpin (residues 43-55) correspond to the previously described “stacking loop” that forms part of the interaction between successive rings of MTP [25](Fig. 2F). The 80α HTJP is quite similar to gp16 of SPP1, with an RMSD of 6.99Å for all equivalent residue pairs, but is more divergent from gpFII of lambda (RMSD=14.43Å), which lacks the extended external loop and has an additional α-helix at its N-terminus that forms a ring around the barrel made from the β-hairpins of gpW [36](Fig. S2; Table S2). The solution structure of gpFII lacks this α-helix [37], suggesting that this ring is formed only when the protein oligomerizes upon binding to the the gpW connector.

### 3. The tail terminator protein

The AlphaFold2 model for gp52 matched the hexameric protein ring below the HTJP (Fig. 1E, 2A-E). By analogy with phage lambda, where it corresponds to gpU, we call this the “tail terminator protein” (TrP) [10,35]. Tail terminators belong to a family of “tail-to-head joining proteins” (THJPs), as defined by Auzat et al. [39], that are added to the tail prior to its attachment to the capsid. Indeed, gp52 was not found in highly purified 80α and SaPI1 procapsids by MS analysis [40], but was present in material that contained 80α tails, consistent with a tail association.

The TrP consists of a four-stranded β-sheet with two α-helices (Fig. 2F), resembling one-half of a TTP fold, indicating a structural and evolutionary relationship, as suggested by Cardarelli et al. [34]. The β-sheet provides a predominantly negatively charged surface that is continuous with the β-sheets of MTP that form the inner lining of the tail tube (Fig 2J). Two loops on one side of the sheet mesh with the extended loops of two HTJP subunits above it, while the loops on the opposite side connects with the MTP [25] (Fig. 2A-D, G-H). The 80α TrP protein is similar to those from lambda (gpU; RMSD=5.4 Å for 114 equivalent atoms) and GTA (Rcc01690; RMSD=10.86 Å) (Fig. S2; Table S2).

Below the TrP, a sixfold symmetric ring could be identified as the major tail protein (gp53), for which the structure was previously determined from the reconstruction of the 80α baseplate [25] (Fig. 1E, 2A-D). It consists of an 8-stranded β-sandwich where one side presents a negatively charged surface to the inside of the tail (Fig. 2F, J). The neck reconstruction showed part of the second MTP ring, but the density was too poor to be modeled. A total of 39 or 40 MTP rings form the complete 80α and SaPI1 tail. In the tail, the C-terminus (C-arm) of MTP interacts with an extended “stacking loop” and extends into the β-sheet of the MTP ring above it [25]. The same C-arm interacts with the shorter stacking loop of TrP, while the N-terminus interacts with the second loop of the adjacent TrP subunit (Fig. 2K).

### 4. The tail interior

In the C6 symmetrical reconstruction, an elongated density could be clearly discerned in the inner tube of the neck stretching all the way from the portal through the neck and into the tail tube; however, the density was not resolvable to high resolution, presumably due to smearing caused by the sixfold averaging. To resolve this internal density, we carried out a symmetry expansion from C6 to C1, followed by masked classification and asymmetric reconstruction focused on the tail interior. The mask was then widened, resulting in a C1 reconstruction encompassing the entire neck that reached a resolution of 3.4 Å (FSC=0.143; Fig. 1D; Fig. S1; Table S1).

This reconstruction clearly showed density corresponding to a double-stranded DNA helix in the upper part of the neck, extending about 72Å from the clip domain of PP, through the HTCP, and ending near the interface between the HTCP and the HTJP (Fig. 1D, F). Immediately below the DNA, tubular densities corresponding to α-helices could be discerned (Fig. 1D, F). We initially expected this density to correspond to the TMP, gp56, which is assumed to extend through the entirety of the tail [41]. To our surprise, these α-helical densities instead matched the AlphaFold2 model of gp51, which could be fitted with high confidence into the density as a monomer (Fig. 3A). To model the internal density accurately, we generated an arbitrary DNA sequence using AlphaFold3, which was combined with the Alphafold2 model of gp51 to initiate the model refinement, resulting in a structure that included residues 7–115 of gp51 and 20 base pairs of B-DNA (Fig. 3A-E).

**Figure 3.**
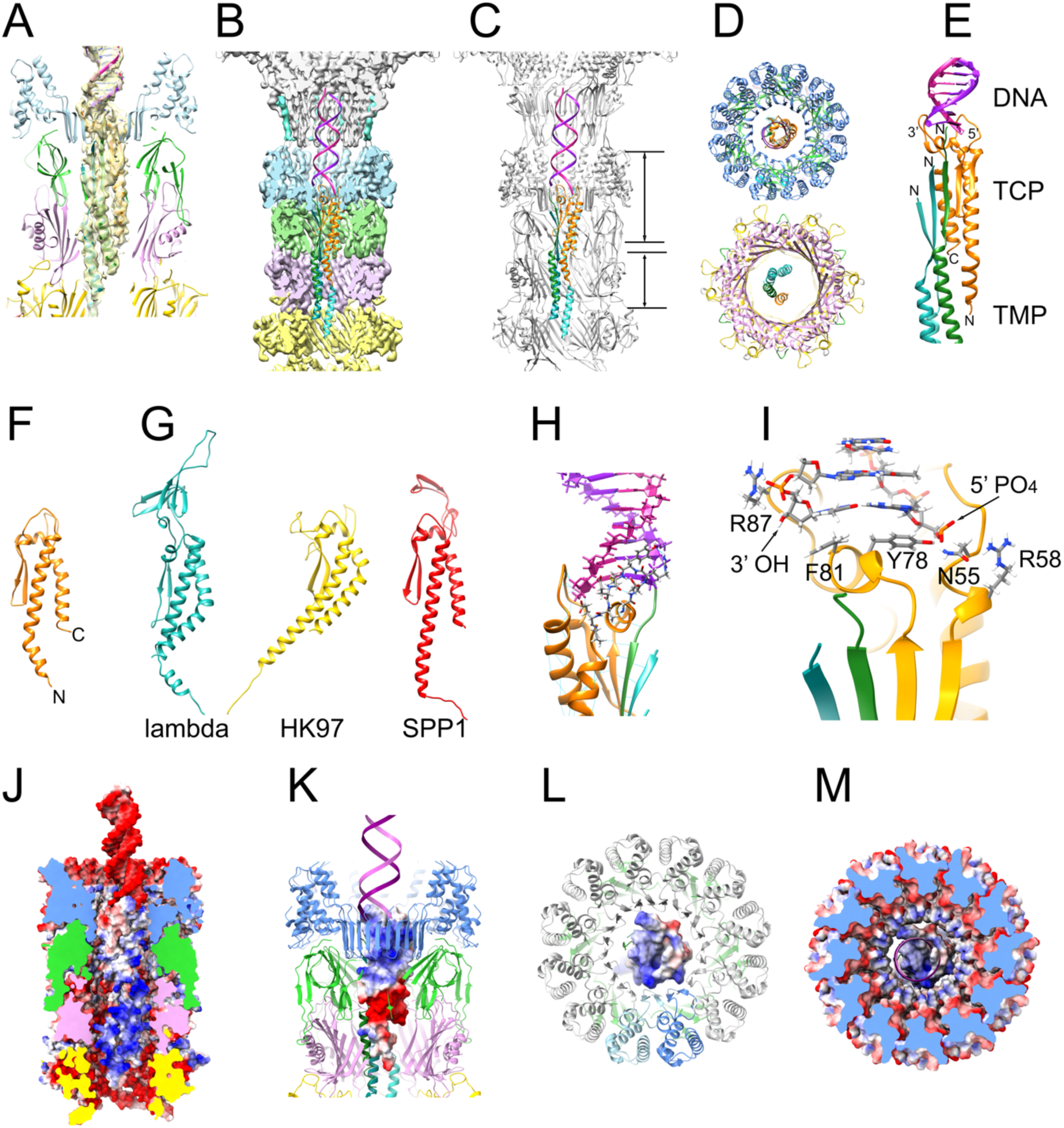
Modeling of the inside of the neck. (A) Detail of internal density with the TCP (gp50, orange), TMP (gp56, shades of green) and DNA (purple) models fitted in. External proteins (HTCP, HTJP, TrP and MTP) are colored as in Fig. 2. (B) Atomic model of the inside of the neck, including the DNA double helix (purple and magenta), TCP (orange) and the three copies of TMP in green, turquoise and aqua, inside the segmented density of the asymmetric reconstruction. (C) Same model as in (B) with the atomic model of the surrounding proteins shown in gray. (D) slabs through the tail model at the levels indicated in (C). (E) Closeup view of the internal proteins and the DNA. (F) Ribbon diagram of the TCP model. (G) Ribbon diagrams of the AlphaFold3 models of the TCPs from phage lambda (gpZ), HK97 (gp10) and SPP1 (gp16.1). (H) The interaction between TCP, TMP and the DNA. The TCP interacting loop is shown in stick representation, colored by element. (I) Detail of the interaction between TCP and the DNA. The 3’ and 5’ ends of the DNA and pertinent residues are indicated. (J) Electrostatic potential surfaces of the inside of the tail. (K) Electrostatic surface of TCP with surrounding proteins and DNA shown in ribbon representation. (L) Top view of the electrostatic surface of TCP with the HTCP shown in ribbon representation. (K) Same view as panel L with TCP and HTCP both shown as electrostatic surfaces.

The gp51 structure consists of a 55 Å long, bent N-terminal α-helix, an extended loop interspersed with a β-hairpin and a short α-helix, followed by a 29 Å long C-terminal α-helix (Fig. 3E, F). Based on the location of ORF51 in the genome (Fig. 1A), gp51 is equivalent to the “tail completion protein” (TCP) of phage lambda (gpZ) [10] and its equivalents gp16.1 in SPP1 [42] and gp10 in HK97. While no structures of these or any other TCPs have been determined, AlphaFold models of lambda gpZ, SPP1 gp16.1 and HK97 gp10 reveal the structural similarity (Fig. 3G). Based on these models, the 80α TCP is most similar to gp16.1 from SPP1 (RMSD=6.93 Å for 108 equivalent Cα atoms), while lambda gp6 is more distantly related, with an additional β domain inserted at its apex (Fig. 3G), perhaps reflecting the difference in packaging strategy. However, the locations of these proteins in their respective phages have not been determined: Lambda gpZ was not observed in recent high-resolution structures of phage lambda [43–45], and although gp16.1 was shown to be tail-associated in SPP1, its exact location and structure has remained unknown [42].

Since there is no unique sequence at the ends of the genome in 80α or SaPI1, it was not possible to model the DNA accurately in our structure. The end of the modeled DNA appeared to be blunt, with a phosphate group at the 5’ end, as expected for DNA packaged by a headful mechanism [29] (Fig. 3A,H,I). However, additional density at the 3’ end of the DNA that could not be modeled might represent a single-nucleotide overhang. Residues Y78 and F81 in the extended loop of the TCP cap the DNA double-helix by forming pi-stacking interactions with the final base pair. (Fig. 3I) The 5’ and first backbone phosphates are stabilized by a hydrophilic pocket formed from residue N55 and exposed backbone peptide bonds in the TCP loop (Fig. 3I). The last backbone phosphate at the 3’ end is stabilized by the guanidinium group of R87 (Fig. 3I). These non-specific interactions accommodate the end of the DNA terminus regardless of DNA sequence.

Below the TCP, several α-helical densities could be discerned, which were presumed to belong to the TMP (Fig. 1D, F, Fig. 3A). Several AlphaFold3 predictions incorporating varying combinations of HTCP, HTJP, TrP, TCP, and DNA, together with three copies of TMP (consistent with our previous reconstruction of the 80α baseplate [25]) were generated. While neither of these models accurately represented the entire neck structure, one model included an interaction between TCP and TMP that accurately matched the density below the TCP. In this model, the N-termini of three copies of TMP each contributes a β-strand to a five-stranded β-sheet that includes the β-hairpin of the TCP (Fig. 3B-E). As a consequence, the α-helices from the three copies of TMP are out of register by about 6-7 residues. The N- and C-terminal α-helices from the TCP together with the three α-helices from TMP thus form a five membered superhelix (Fig. 3D-E). Only residues 1–37 of TMP could be modeled into the density. Below this, the density deteriorates and the organization of TMP becomes unclear (Fig. 1D). Once its C-terminus reaches the baseplate, however, the three copies must be back into register again, since the three C-termini are known to interact with the Tal protein with identical interactions, as shown in our previous reconstruction of the 80α baseplate [25]. It is not clear how this is accomplished, but TMP is often seen by SDS-PAGE to be fragmented, suggesting that it might be cleaved in some way inside the tail.

The TMP is predominantly positively charged, complementary to the negatively charged inner lining of the neck and tail (Fig. 3J), presumably providing resistance against the pressure from the packaged DNA. The TCP forms a connector between the DNA and the TMP, presenting a positively charged surface to the DNA and a negatively charged surface to the TMP (Fig. 3K-M).

There are minimal interactions between the interior neck proteins (TCP and TMP) and either of the exterior proteins (HTCP, HTJP, TrP and MTP). Interactions between TCP and the HTCP dodecamer make up 60% of the total buried surface area between the interior and exterior neck proteins, about half (48%) of which involves two adjacent HTCP monomers that interface with a large positively charged surface at the base of the TCP extended loop (Fig. 3K-M), suggesting that this area is primarily responsible for stabilizing the position of the TCP, and thereby the DNA and TMP, in the neck. The HTCPs from HK97 and JBD30 lack the β-hairpin that interacts with TCP in 80α (Fig. S1). It is unclear if the TCPs of these phages are located inside the neck structure and how they might interact with the HTCP [32].

## DISCUSSION

Here, we have determined the structure of the neck of SaPI1 virions, consisting of structural proteins provided by the 80α helper phage. We were able to resolve all the proteins that form the connecting region between the portal and the major tail protein, as well as the DNA and proteins on the inside of the tail. We have also resolved a long-standing question in the field: the structure, location and interaction of the tail completion and tape measure proteins.

The terminology used to describe the proteins involved in the head-to-tail connection in various systems is somewhat confusing. In some systems, such as ϕ29 and P2, the portal is traditionally called the connector, though the term “connector” (or “head-to-tail connector”) is more often associated with the protein ring below the portal and sometimes to the whole complex of proteins between head and tail. We have chosen to refer to the protein making up the ring immediately below the portal as the head-tail connector protein (HTCP), and the ring below it as the head-tail joining protein (HTJP), consistent with usage in phage lambda (Table 1). In SPP1 the two proteins are referred to as “adaptor” and “stopper”, but this implies a functional role that has not been fully characterized in 80α. This organization with two “connector” rings (HTCP and HTJP) is common, but not universal: in the T5-like *E. coli* phage DT57C, there is no HTJP equivalent; instead, the HTCP appears to be highly modified to interact directly with the TrP [46]. The same might be true in JBD30, where the protein identified as “stopper” is, in fact, a TrP-like protein [32]. It has been proposed that the types of HTCP present in lambda (gpW) and HK97 (gp6) are unrelated [35], but our analysis suggests that the two folds are in fact structurally and presumably evolutionarily related (Fig. S1). The HTCP and HTJP proteins are also referred to collectively as “head completion proteins” to reflect the fact that they are added to the heads after assembly, but before joining with the tail [10,34].

Similarly, “tail completion proteins” are proteins added to the tail prior to head-tail joining. This includes the tail terminator (TrP), which caps off the tail after addition of a sufficient number of rings of MTP [41]. However, “tail completion protein” (TCP) is also used to more specifically denote lambda protein gpZ and its homologs [10,35], which includes 80α gp51, identified in this study. Until the present study, the structure and location of these ubiquitous proteins was unknown. In the 80α/SaPI1 tail, its location inside the neck suggests that it might be involved in both head-tail joining and DNA ejection, consistent with the observations in SPP1 [42]. To avoid confusion, we have chosen to retain the “tail completion protein” (TCP) terminology for gp51.

DNA packaging requires the terminase complex, consisting of TerS and TerL, which dock with the portal protein in the procapsids to initiate packaging. TerL, at least, must remain attached to the portal during packaging, presumably via the clip domain of the PP [30]. Since TerL is presumed to be a pentamer [47], there is a symmetry mismatch between the portal and the terminase. Upon completion of packaging, TerL is removed and is replaced by the HTCP, which, like the portal, is a dodecamer and forms a very tight interaction with the PP [30]. How this exchange between TerL and HTCP is coordinated in order to avoid DNA escape is not clear. Presumably, the DNA remains inside the portal, perhaps involving the channel loops of the PP [30], at least until the HTCP ring is added. In phage lambda, the HTCP itself, comprised of gpW, provides a narrow channel (20 Å wide) that might be sufficient to prevent DNA leakage [43,45]. In the 80α HTCP, this channel is 31 Å at its narrowest (Fig. 2E), probably too wide to prevent DNA escape. Instead, the hexameric HTJP ring might serve in this role. In SPP1, gp16 residues 40-60 in neck complexes extracted from tail-less, DNA-filled capsids formed a ring of α-helices that narrowed the diameter of the internal tunnel of the HTJP ring to ≈11Å and was thus proposed to act as a “stopper” [31]. Although the 80α HTJP lacks this α-helix, and the HTJP tunnel is ≈36 Å wide at its narrowest, the AlphaFold2 model for gp50 predicted an α-helix in residues 43–52 (Fig. 2I), suggesting that the open state we observed might represent a reorganized protein that forms once the HTJP ring interacts with the tail (the ring formed by TrP). Once the tail is attached, the DNA would be blocked by the TCP, which occupies the central tunnel.

Long-tailed phages (siphoviruses and myoviruses) assemble their tails through a pathway separate from that of the head, followed by attachment to the heads after completion of DNA packaging. The TMP is a large protein that defines the length of the tail [41]. By comparison with other systems, tail assembly is assumed to be nucleated from a complex made from Tal, TMP and the distal tail protein (Dit) and proceed by addition of hexameric MTP rings (39 or 40 rings in the case of 80α and SaPI1) until it reaches the end of the fully extended TMP, upon which the terminator TrP is added to cap off the stack of MTP rings [41]. In phage lambda, the TCP (gpZ) is the last protein to be added to the tail prior to head-tail attachment [41]. It was proposed that gpZ binds specifically to one end of the DNA [48]. This may differ from 80α and other headful packaging phages, where the packaged DNA does not have a specific sequence at the ends. The location of gpZ is not known and it was not observed in high-resolution reconstructions of lambda virions [43,45]. Nor was the TCP observed in high-resolution reconstructions of JBD30 or DT57C [32,46].

Here, we have shown that a monomer of TCP (gp51) is located inside the 80α tail where it interacts with the N-termini of the TMP trimer in an asymmetric manner. The tight five stranded β-sheet formed between TMP and TCP might suggest that the complex is formed early in tail assembly, but there is no additional evidence to support this conjecture. Indeed, gp51 was not seen by MS in a crude procapsid preparation, which includes numerous tails and tail proteins, suggesting that the TCP is either added late in tail assembly, during head-tail joining, or only after DNA packaging [40]. Its location inside the HTJP—assumed to be a part of the head—also points to the TCP being the last protein to be added to the tail, maybe during head-tail joining, at which point it may serve to plug up the tail and prevent the DNA from spontaneously ejecting (functionally a “stopper” protein).

Consistent with this role, absence of gp16.1 in SPP1 led to a 100-fold reduction in titer due to a failure to join heads and tails, and also affected injection of DNA through the cell wall [42]. Similarly, in phage lambda, complete virions could be assembled in the absence of gpZ, but had 500-fold lower infectivity [48]. The location of neither gpZ nor SPP1 gp16.1 has been ascertained, but in light of the similarity of their predicted structures to the 80α TCP, it seems likely that they could occupy a similar location inside their respective tails.

In phage lambda, the DNA was observed far inside the tail, suggesting that it drops down during head-tail joining, ready for ejection [43]. Likewise, in SPP1, between 55 and 67 base pairs of genomic DNA were protected by the tail [49]. In contrast, in the SaPI1/80α tail, the DNA also extends only as far down as the barrel of β-hairpins in the HTCP, where it is blocked from sliding further down by the TCP (Fig. 3). The same is the case in JBD30 and DT57C [32,46]. Thus, either the HTCP itself is capable of holding back the DNA until head-tail joining occurs, or the TCP would have to push the DNA back into the head, perhaps using the TMP as a spring.

In addition to its role in tail assembly and length determination, the TMP is also involved in the DNA ejection process, probably being ejected along with the DNA as a kind of “pilot” protein [25]. We previously showed how the C-terminal α-helices of the TMPs interact with the α-helices in the Tal protein at the tip of the baseplate, providing a model for how interaction of Tal with the host cell membrane could cause a conformational change that is coupled to TMP release to initiate DNA ejection [25]. Given the location of TCP inside the tail, it too presumably escapes together with the TMP and the DNA, consistent with the observed effect on DNA routing in SPP1 [42]. It is worth noting that the 80α minor capsid protein gp44, which serves to protect the DNA from degradation post injection, is also ejected together with the DNA [50,51]. It is not known whether this protein has a specific location in the capsid or the tail or how it might be ejected along with the DNA. We could not have observed gp44 in this study, because the SaPI1 construct that we used was made from an 80α Δ44 deletion, which we previously showed to slightly increase SaPI1 yields, while greatly reducing phage titers [27].

The reconstruction of the SaPI1 neck presented here is the first structure of the neck region of a staphylococcal siphovirus, and the first detailed description of the organization of the DNA, TCP and TMP in any phage. It seems likely that many other long-tailed phages, both siphoviruses and myoviruses, may share the same structure and organization of the TCP in the tail. In some cases, the failure to observe the TCP in high resolution reconstruction of phage necks and tails may be due to the application of threefold averaging during the reconstruction procedure [46]. It could also be that the TCP does not always have a consistent orientation relative to the outer neck proteins in all phages: indeed, in the 80α tail, the TCP is stabilized by minimal interactions with the HTCP. Additionally, phages undergoing sequence-specific DNA packaging (such as lambda) may employ other strategies for positioning their DNA in the tail and readying it for ejection.

## MATERIALS AND METHODS

### 1. Production of SaPI1 virions

The production and purification of SaPI1 virions were described previously [30]. Briefly, SaPI1 virions were produced by mitomycin C induction of *S. aureus* strain ST65, which contains SaPI1 *tst::tetM* as well as an 80α prophage with a deletion of ORF44 (80αΔ*44*) [27]. The virions were purified by PEG 6,000 precipitation, followed by CsCl and sucrose gradient centrifugation. Fractions containing SaPI1 particles were concentrated by pelleting and resuspended in phage dialysis buffer (20 mM Tris-HCl pH 7.8, 50 mM NaCl, 4 mM CaCl_2_, 1 mM MgSO_4_).

### 2. Electron microscopy

Cryo-EM was carried out as previously described [30]. SaPI1 particles were treated with 1µl Benzonase® nuclease at room temperature for 1 hour prior to cryo-EM grid preparation, followed by dialysis on a 0.025 µm MCE Membrane filter (MF-Millipore) floating on phage dialysis buffer. Cryo-EM samples were prepared using a Vitrobot Mark IV with glow discharged nickel Quantifoil R2/1 grids, and imaged using an FEI Titan Krios microscope operated at 300 kV and equipped with a Gatan K3 detector and Quantum GIF energy filter at the Midwestern Center for Cryo-Electron Microscopy (MCCEM) at Purdue University. A total of 2,796 images were collected at a magnification of 64,000 x (pixel size 1.33 Å), and an electron dose of 35.26 e^-^/Å^2^ (Table S1).

### 3. Three-dimensional reconstruction of the SaPI1 neck

Reconstruction of the neck was done in cryoSPARC [52]. The previously picked particle images (59,457 particles) of the full capsids [30] were re-centered on the neck region and reconstructed with the application of C6 symmetry, reaching a final resolution of 3.1Å (FSC=0.143) (Fig. S1; Table S1). To resolve the internal structures, the particles from the C6 reconstruction were symmetry expanded to C1 (yielding a total of 356,742 particles). A mask was applied that only covered the internal density, followed by 3D classification without alignment with 10 classes. The class showing the most clearly resolved internal density (53,086 particles) was refined with a static mask after removing duplicates (final 35,724 particles). Subsequently, the mask was expanded to encompass the whole neck region and refined without symmetry. The final C1 reconstruction reached a resolution of 3.54 Å (Fig. S1; Table S1).

### 4. Model building

Initial models were made in AlphaFold2 [33] and AlphaFold3 [53]. Model building was done in ChimeraX [54] with refinement using ISOLDE [55]. RMSDs were calculated from within UCSF Chimera [56] using the BLOSUM-62 matrix, a pruning distance of 3.5Å, and a secondary structure weight of 50%.

## Data availability

The final maps and coordinates were submitted to EMdep with identifiers EMD-XXXX and PDB IDs YYYY.

## Acknowledgements

We are grateful to Drs. Thomas Klose and Xueyong Xu at the Midwestern Center for Cryo-Electron Microscopy (MCCEM) at Purdue University for assistance with collecting the cryo-EM data. MCCEM was supported by NIH grant U24 GM116789 to Dr. Wen Jiang at Purdue University. Data collection and processing was done with assistance from the UAB Cryo-EM Facility (CEMF), supported by the UAB Institutional Research Core Program (IRCP), the O’Neal Comprehensive Cancer Center (NIH grant P30 CA013148) and NIH grant S10 OD024978 to T.D. This work was supported by NIH research grant R01 AI083255 to T.D.

## SUPPLEMENTARY FIGURE LEGENDS

**Figure S1.** Fourier Shell Correlation (FSC) curves from cryoSPARC for the C6 (A) and C1 (B) reconstructions.

**Figure S2.** Comparison of 80α neck proteins with the equivalent proteins from other phages. (A) HTCP: 80α gp49 (blue), SPP1 gp15 (red), Lambda gpW (purple), HK97 gp6 (yellow), JBD30 gp41 (green), GTA Rcc01688 (tan). (B) HTJP: 80α gp50 (green), SPP1 gp16 (red), lambda gpFII (blue), GTA Rcc01689 (tan). (C) TrP: 80α gp52 (pink), lambda gpU (blue), GTA Rcc01690 (tan).

## REFERENCES

1. Puxty RJ, Millard AD (2023) Functional ecology of bacteriophages in the environment. Curr Opin Microbiol 71: 102245.

2. Taylor VL, Fitzpatrick AD, Islam Z, Maxwell KL (2019) The Diverse Impacts of Phage Morons on Bacterial Fitness and Virulence. Adv Virus Res 103: 1–31.

3. Schroven K, Aertsen A, Lavigne R (2021) Bacteriophages as drivers of bacterial virulence and their potential for biotechnological exploitation. FEMS Microbiol Rev 45: fuaa041.

4. Rao VB, Fokine A, Fang Q (2021) The remarkable viral portal vertex: structure and a plausible model for mechanism. Curr Opin Virol 51: 65–73.

5. Dedeo CL, Cingolani G, Teschke CM (2019) Portal Protein: The Orchestrator of Capsid Assembly for the dsDNA Tailed Bacteriophages and Herpesviruses. Annu Rev Virol 6: 141–160.

6. Prevelige PE, Cortines JR (2018) Phage assembly and the special role of the portal protein. Curr Opin Virol 31: 66–73.

7. Turner D, Shkoporov AN, Lood C, Millard AD, Dutilh BE, Alfenas-Zerbini P, van Zyl LJ, Aziz RK, Oksanen HM, Poranen MM, Kropinski AM, Barylski J, Brister JR, Chanisvili N, Edwards RA, Enault F, Gillis A, Knezevic P, Krupovic M, Kurtböke I, Kushkina A, Lavigne R, Lehman S, Lobocka M, Moraru C, Moreno Switt A, Morozova V, Nakavuma J, Reyes Muñoz A, Rūmnieks J, Sarkar BL, Sullivan MB, Uchiyama J, Wittmann J, Yigang T, Adriaenssens EM (2023) Abolishment of morphology-based taxa and change to binomial species names: 2022 taxonomy update of the ICTV bacterial viruses subcommittee. Arch Virol 168: 74.

8. Veesler D, Cambillau C (2011) A common evolutionary origin for tailed-bacteriophage functional modules and bacterial machineries. Microbiol Mol Biol Rev 75: 423–433.

9. Leiman PG, Shneider MM (2012) Contractile tail machines of bacteriophages. Adv Exp Med Biol 726: 93–114.

10. Davidson AR, Cardarelli L, Pell LG, Radford DR, Maxwell KL (2012) Long noncontractile tail machines of bacteriophages. Adv Exp Med Biol 726: 115–142.

11. Tavares P (2018) The Bacteriophage Head-to-Tail Interface. Subcell Biochem 88: 305–328.

12. Kourtis AP, Hatfield K, Baggs J, Mu Y, See I, Epson E, Nadle J, Kainer MA, Dumyati G, Petit S, Ray SM, Emerging IPMRSAAG, Ham D, Capers C, Ewing H, Coffin N, McDonald LC, Jernigan J, Cardo D (2019) Vital Signs: Epidemiology and Recent Trends in Methicillin-Resistant and in Methicillin-Susceptible Staphylococcus aureus Bloodstream Infections - United States. MMWR Morb Mortal Wkly Rep 68: 214–219.

13. Archer GL (1998) Staphylococcus aureus: A well-armed pathogen. Clin Infect Dis 26: 1179–1181.

14. Malachowa N, DeLeo FR (2010) Mobile genetic elements of Staphylococcus aureus. Cell Mol Life Sci 67: 3057–3071.

15. Lindsay JA (2014) Staphylococcus aureus genomics and the impact of horizontal gene transfer. Int J Med Microbiol 304: 103–109.

16. Xia G, Wolz C (2014) Phages of Staphylococcus aureus and their impact on host evolution. Infect Genet Evol 21: 593–601.

17. Chiang YN, Penadés JR, Chen J (2019) Genetic transduction by phages and chromosomal islands: The new and noncanonical. PLoS Pathog 15: e1007878.

18. Novick RP, Christie GE, Penades JR (2010) The phage-related chromosomal islands of Gram-positive bacteria. Nat Rev Microbiol 8: 541–551.

19. Penadés JR, Christie GE (2015) The Phage-Inducible Chromosomal Islands: A Family of Highly Evolved Molecular Parasites. Annu Rev Virol 2: 181–201.

20. Christie GE, Dokland T (2012) Pirates of the Caudovirales. Virology 434: 210–221.

21. Dokland T (2019) Molecular Piracy: Redirection of Bacteriophage Capsid Assembly by Mobile Genetic Elements. Viruses 11: 1003.

22. Diep BA, Gill SR, Chang RF, Phan TH, Chen JH, Davidson MG, Lin F, Lin J, Carleton HA, Mongodin EF, Sensabaugh GF, Perdreau-Remington F (2006) Complete genome sequence of USA300, an epidemic clone of community-acquired meticillin-resistant Staphylococcus aureus. Lancet 367: 731–739.

23. Spilman MS, Dearborn AD, Chang JR, Damle PK, Christie GE, Dokland T (2011) A conformational switch involved in maturation of Staphylococcus aureus bacteriophage 80alpha capsids. J Mol Biol 405: 863–876.

24. Kizziah JL, Manning KA, Dearborn AD, Wall EA, Klenow L, Hill RLL, Spilman MS, Stagg SM, Christie GE, Dokland T (2017) Cleavage and Structural Transitions during Maturation of Staphylococcus aureus Bacteriophage 80α and SaPI1 Capsids. Viruses 9: E384.

25. Kizziah JL, Manning KA, Dearborn AD, Dokland T (2020) Structure of the host cell recognition and penetration machinery of a Staphylococcus aureus bacteriophage. PLoS Pathog 16: e1008314.

26. Christie GE, Matthews AM, King DG, Lane KD, Olivarez NP, Tallent SM, Gill SR, Novick RP (2010) The complete genomes of Staphylococcus aureus bacteriophages 80 and 80 alpha - implications for the specificity of SaPI mobilization. Virology 407: 381–390.

27. Dearborn AD, Spilman MS, Damle PK, Chang JR, Monroe EB, Saad JS, Christie GE, Dokland T (2011) The Staphylococcus aureus pathogenicity island protein gp6 functions as an internal scaffold during capsid size determination. J Mol Biol 412: 710–722.

28. Dearborn AD, Wall EA, Kizziah JL, Klenow L, Parker LK, Manning KA, Spilman MS, Spear JM, Christie GE, Dokland T (2017) Competing scaffolding proteins determine capsid size during mobilization of Staphylococcus aureus pathogenicity islands. Elife 6: 10.7554.

29. Casjens SR, Gilcrease EB (2009) Determining DNA packaging strategy by analysis of the termini of the chromosomes in tailed-bacteriophage virions. Methods Mol Biol 502: 91–111.

30. Mukherjee A, Kizziah JL, Hawkins NC, Nasef MO, Parker LK, Dokland T (2024) Structure of the Portal Complex from Staphylococcus aureus Pathogenicity Island 1 Transducing Particles In Situ and In Isolation. J Mol Biol 436: 168415.

31. Orlov I, Roche S, Brasilès S, Lukoyanova N, Vaney MC, Tavares P, Orlova EV (2022) CryoEM structure and assembly mechanism of a bacterial virus genome gatekeeper. Nat Commun 13: 7283.

32. Valentová L, Füzik T, Nováček J, Hlavenková Z, Pospíšil J, Plevka P (2024) Structure and replication of Pseudomonas aeruginosa phage JBD30. EMBO J 43: 4384–4405.

33. Jumper J, Evans R, Pritzel A, Green T, Figurnov M, Ronneberger O, Tunyasuvunakool K, Bates R, Žídek A, Potapenko A, Bridgland A, Meyer C, Kohl SAA, Ballard AJ, Cowie A, Romera-Paredes B, Nikolov S, Jain R, Adler J, Back T, Petersen S, Reiman D, Clancy E, Zielinski M, Steinegger M, Pacholska M, Berghammer T, Bodenstein S, Silver D, Vinyals O, Senior AW, Kavukcuoglu K, Kohli P, Hassabis D (2021) Highly accurate protein structure prediction with AlphaFold. Nature 596: 583–589.

34. Cardarelli L, Lam R, Tuite A, Baker LA, Sadowski PD, Radford DR, Rubinstein JL, Battaile KP, Chirgadze N, Maxwell KL, Davidson AR (2010) The crystal structure of bacteriophage HK97 gp6: defining a large family of head-tail connector proteins. J Mol Biol 395: 754–768.

35. Casjens SR, Davidson AR, Grose JH (2022) The small genome, virulent, non-contractile tailed bacteriophages that infect Enterobacteriales hosts. Virology 573: 151–166.

36. Maxwell KL, Yee AA, Booth V, Arrowsmith CH, Gold M, Davidson AR (2001) The solution structure of bacteriophage lambda protein W, a small morphogenetic protein possessing a novel fold. J Mol Biol 308: 9–14.

37. Maxwell KL, Yee AA, Arrowsmith CH, Gold M, Davidson AR (2002) The solution structure of the bacteriophage lambda head-tail joining protein, gpFII. J Mol Biol 318: 1395–1404.

38. Bárdy P, Füzik T, Hrebík D, Pantůček R, Thomas Beatty J, Plevka P (2020) Structure and mechanism of DNA delivery of a gene transfer agent. Nat Commun 11: 3034.

39. Auzat I, Petitpas I, Lurz R, Weise F, Tavares P (2014) A touch of glue to complete bacteriophage assembly: the tail-to-head joining protein (THJP) family. Mol Microbiol 91: 1164–1178.

40. Poliakov A, Chang JR, Spilman MS, Damle PK, Christie GE, Mobley JA, Dokland T (2008) Capsid size determination by *Staphylococcus aureus* pathogenicity island SaPI1 involves specific incorporation of SaPI1 proteins into procapsids. J Mol Biol 380: 465–475.

41. Katsura I (1990) Mechanism of length determination in bacteriophage lambda tails. Adv Biophys 26: 1–18.

42. Auzat I, Ouldali M, Jacquet E, Fauler B, Mielke T, Tavares P (2024) Dual function of a highly conserved bacteriophage tail completion protein essential for bacteriophage infectivity. Commun Biol 7: 590.

43. Gu Z, Wu K, Wang J (2024) Structural morphing in the viral portal vertex of bacteriophage lambda. J Virol 98: e0006824.

44. Wang C, Duan J, Gu Z, Ge X, Zeng J, Wang J (2024) Architecture of the bacteriophage lambda tail. Structure 32: 35–46.e3.

45. Xiao H, Tan L, Tan Z, Zhang Y, Chen W, Li X, Song J, Cheng L, Liu H (2023) Structure of the siphophage neck-Tail complex suggests that conserved tail tip proteins facilitate receptor binding and tail assembly. PLoS Biol 21: e3002441.

46. Ayala R, Moiseenko AV, Chen TH, Kulikov EE, Golomidova AK, Orekhov PS, Street MA, Sokolova OS, Letarov AV, Wolf M (2023) Nearly complete structure of bacteriophage DT57C reveals architecture of head-to-tail interface and lateral tail fibers. Nat Commun 14: 8205.

47. Hawkins DEDP, Bayfield OW, Fung HKH, Grba DN, Huet A, Conway JF, Antson AA (2023) Insights into a viral motor: the structure of the HK97 packaging termination assembly. Nucleic Acids Res 51: 7025–7035.

48. Thomas JO, Sternberg N, Weisberg R (1978) Altered arrangement of the DNA in injection-defective lambda bacteriophage. J Mol Biol 123: 149–161.

49. Tavares P, Lurz R, Stiege A, Rückert B, Trautner TA (1996) Sequential headful packaging and fate of the cleaved DNA ends in bacteriophage SPP1. J Mol Biol 264: 954–967.

50. Manning KA, Quiles-Puchalt N, Penadés JR, Dokland T (2018) A novel ejection protein from bacteriophage 80α that promotes lytic growth. Virology 525: 237–247.

51. Manning KA, Dokland T (2020) The gp44 Ejection Protein of Staphylococcus aureus Bacteriophage 80α Binds to the Ends of the Genome and Protects It from Degradation. Viruses 12: 563.

52. Punjani A, Rubinstein JL, Fleet DJ, Brubaker MA (2017) cryoSPARC: algorithms for rapid unsupervised cryo-EM structure determination. Nat Methods 14: 290–296.

53. Roy R, Al-Hashimi HM (2024) AlphaFold3 takes a step toward decoding molecular behavior and biological computation. Nat Struct Mol Biol 31: 997–1000.

54. Pettersen EF, Goddard TD, Huang CC, Meng EC, Couch GS, Croll TI, Morris JH, Ferrin TE (2021) UCSF ChimeraX: Structure visualization for researchers, educators, and developers. Protein Sci 30: 70–82.

55. Croll TI (2018) ISOLDE: a physically realistic environment for model building into low-resolution electron-density maps. Acta Crystallogr D Struct Biol 74: 519–530.

56. Goddard TD, Huang CC, Ferrin TE (2007) Visualizing density maps with UCSF Chimera. J Struct Biol 157: 281–287.

57. Cardarelli L, Pell LG, Neudecker P, Pirani N, Liu A, Baker LA, Rubinstein JL, Maxwell KL, Davidson AR (2010) Phages have adapted the same protein fold to fulfill multiple functions in virion assembly. Proc Natl Acad Sci U S A 107: 14384–14389.

58. Huet A, Oh B, Maurer J, Duda RL, Conway JF (2023) A symmetry mismatch unraveled: How phage HK97 scaffold flexibly accommodates a 12-fold pore at a 5-fold viral capsid vertex. Sci Adv 9: eadg8868.

59. Juhala RJ, Ford ME, Duda RL, Youlton A, Hatfull GF, Hendrix RW (2000) Genomic sequences of bacteriophages HK97 and HK022: pervasive genetic mosaicism in the lambdoid bacteriophages. J Mol Biol 299: 27–51.

