## Supplemental Figures and Tables for "Structure of the *Staphylococcus aureus* bacteriophage 80α neck shows the interactions between DNA, tail completion protein and tape measure protein"

### SUPPLEMENTARY FIGURE LEGENDS

**Figure S1.** Fourier Shell Correlation (FSC) curves from cryoSPARC for the C6 (A) and C1 (B) reconstructions.

**Figure S2.** Comparison of 80$\alpha$ neck proteins with the equivalent proteins from other phages. (A) HTCP: 80$\alpha$ gp49 (blue), SPP1 gp15 (red), Lambda gpW (purple), HK97 gp6 (yellow), JBD30 gp41 (green), GTA Rcc01688 (tan). (B) HTJP: 80$\alpha$ gp50 (green), SPP1 gp16 (red), lambda gpFII (blue), GTA Rcc01689 (tan). (C) TrP: 80$\alpha$ gp52 (pink), lambda gpU (blue), GTA Rcc01690 (tan).

**SUPPLEMENTARY FIGURES**

**Figure S1**

A

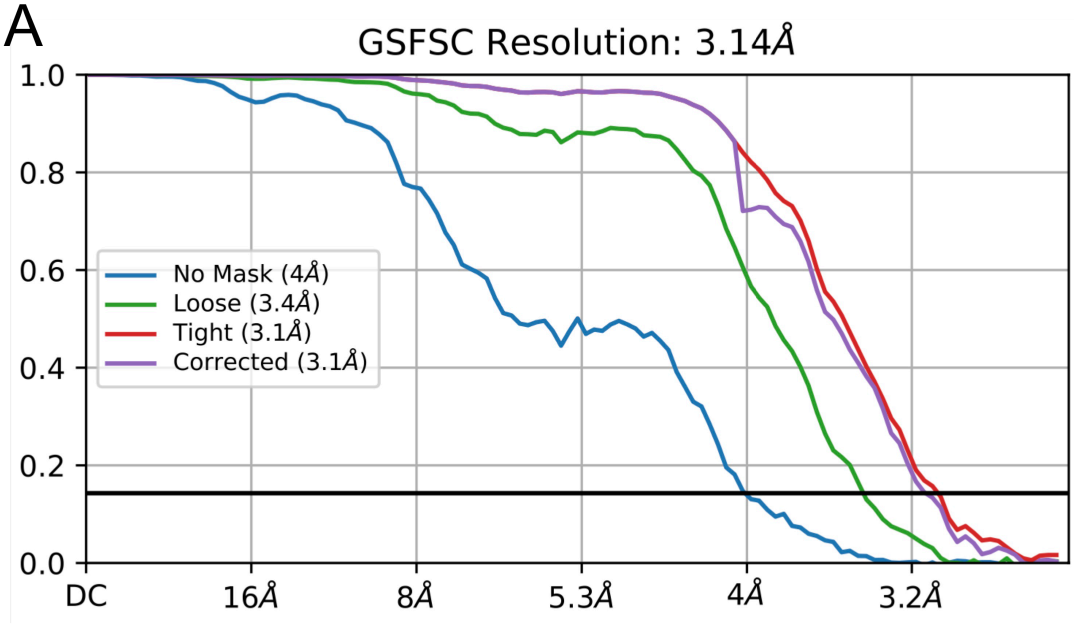


B

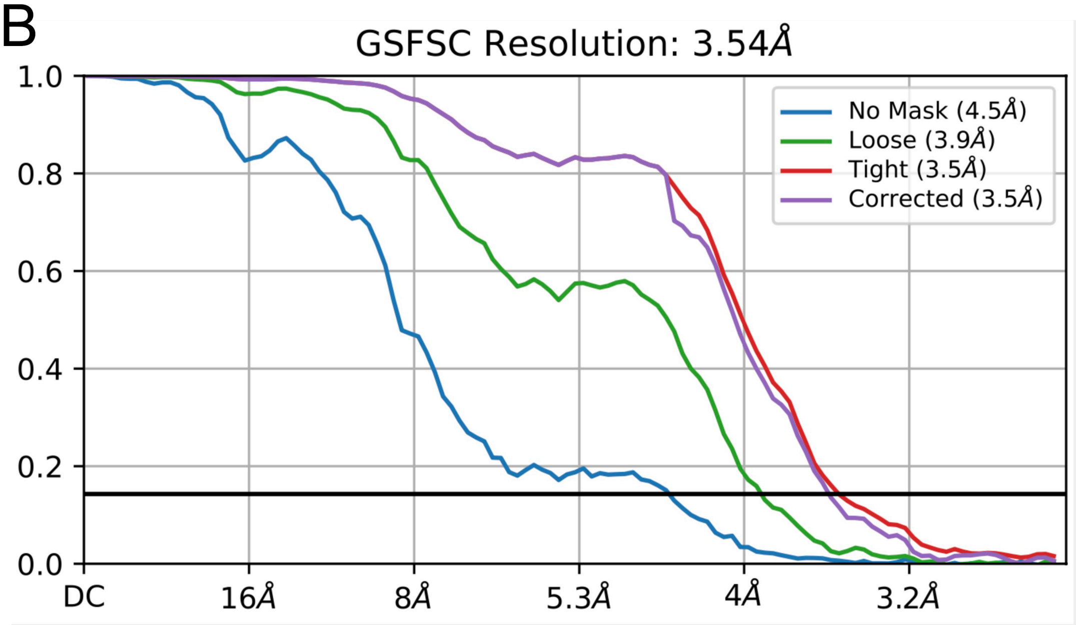

**Figure S2**

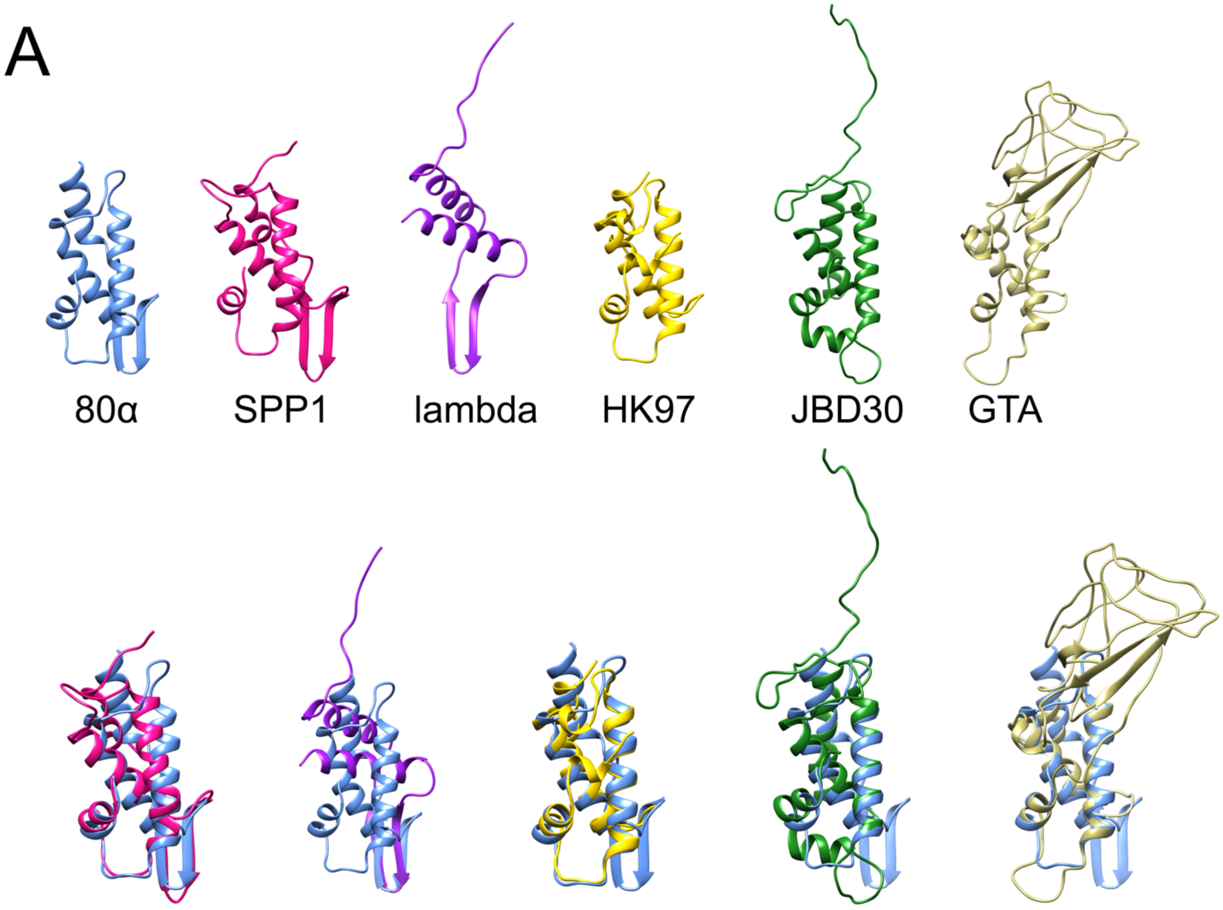

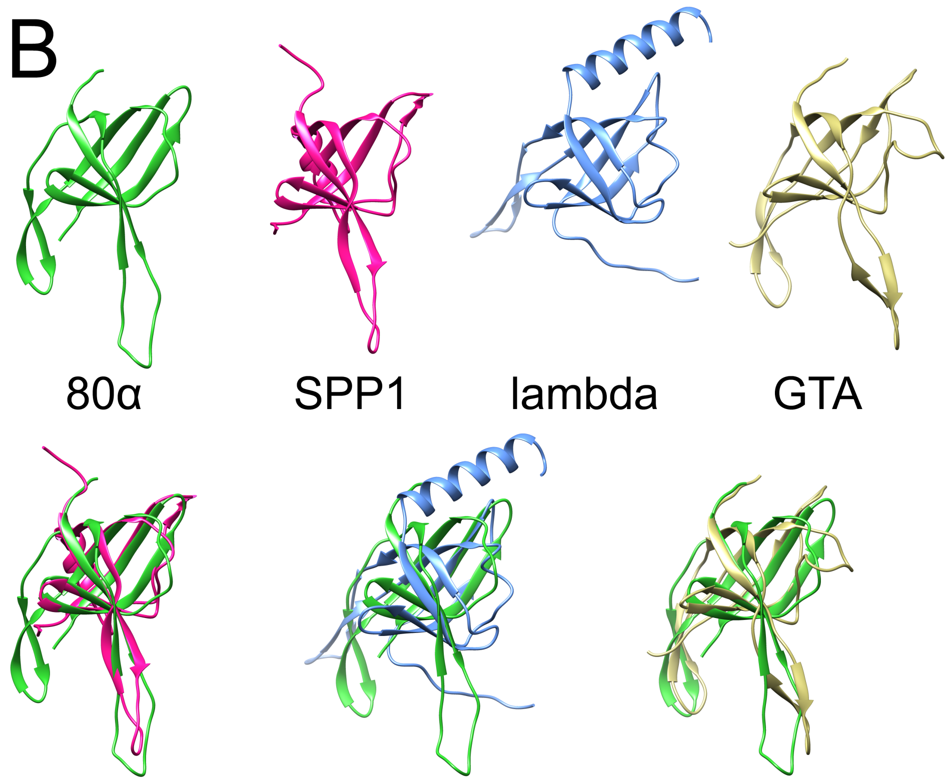
B
80α
SPP1
lambda
GTA

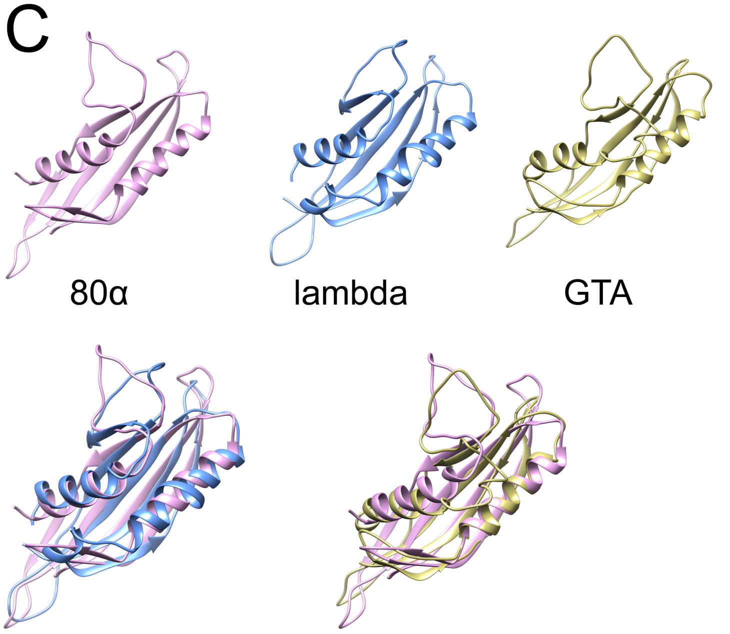
C
80α
lambda
GTA

### SUPPLEMENTARY TABLES

**Supplementary Table S1. (A)** Data collection and processing parameters and statistics.

| Structure | Neck C6 | Neck C1 |
|---|---|---|
| Microscope | FEI Titan Krios G1 | |
| Camera | Gatan K3 | |
| Energy filter | Gatan Quantum GIF | |
| Image collection software | Leginon | |
| Slit width (eV) | 20 | |
| Voltage (kV) | 300 | |
| Micrographs collected | 2,796 | |
| Defocus range (µm) | 0.7–2.0 | |
| Total exposure ($e^-$/Å$^2$) | 35.26 | |
| Frames per movie | 38 | |
| Pixel size (Å/pix) | 1.33 | |
| Reconstruction software | RELION-4.0; CryoSPARC v4.2 | |
| Final particles | 59,457 | 35,724 |
| Symmetry imposed | C6 | C1 |
| Map resolution (Å, $FSC_{0.143}$) | 3.1 | 3.5 |
| EMDB Accession | EMD-XXXX | EMD-YYYY |

**Supplementary Table S1. (B)** Model building parameters and statistics.

| **Structure** | **Neck C6** | **Neck C1** |
|---|---|---|
| Refinement resolution (Å) | 3.1 | 3.5 |
| Model composition: | | |
| Chains | 6 | 42 |
| Atoms | 4,822 | 31,584 |
| Hydrogens | 0 | 0 |
| Protein residues | 593 | 3,779 |
| Nucleotides | 0 | 40 |
| Waters | 0 | 0 |
| Ligands | 0 | 0 |
| Bonds (RMSD): | | |
| Length (Å) (# > 4σ) | 0.006 (0) | 0.006 (0) |
| Angles (°) (# > 4σ) | 1.230 (0) | 1.152 (0) |
| MolProbity score | 0.69 | 0.88 |
| Clash score | 0.00 | 0.52 |
| Ramachandran plot (%): | | |
| Favored | 69.90 | 96.73 |
| Allowed | 3.10 | 3.27 |
| Outliers | 0.00 | 0.00 |
| Rama-Z score (RMSD), N: | | |
| Whole | -0.45 (0.33), N=581 | -0.69 (0.13), N=3,699 |
| Helix | -1.46 (0.35), N=153 | -1.34 (0.13), N=1,055 |
| Sheet | 0.41 (0.37), N=185 | 0.24 (0.15), N=1,212 |
| Loop | 0.16 (0.40), N=243 | 0.03 (0.16), N=1,432 |
| Rotamer outliers (%) | 0.0 | 0.0 |
| $C_{\beta}$ outliers (%) | 0.0 | 0.0 |
| Peptide plane (%): | | |
| Cis proline/general | 12.5/0.0 | 12.2/0.0 |
| Twisted proline/general | 0.0/0.0 | 0.0/0.0 |
| CaBLAM outliers (%) | 0.35 | 0.50 |
| Model fit vs. Map: | | |
| $FSC_{0.5}$ ($FSC_{0.143}$) | 3.4 (3.1) | 3.7 (3.5) |
| $CC_{volume}$ | 0.80 | 0.83 |
| $CC_{mask}$ | 0.86 | 0.83 |
| Accession (PDB) | XXXX | YYYY |

**Supplementary Table S2.** Root-mean-square deviations (RMSD) of equivalent Cα atoms between 80α/SaPI1 neck proteins and the equivalent proteins from other bacteriophages.

| **80α/SaPI1** | | **vs.** | **SPP1** | **PDB ID: 7Z4W** | | | |
|---|---|---|---|---|---|---|---|
| Protein | Gene product | # res | gp | Total¶ | RMSD | Pruned§ | RMSD |
| HTCP | gp49 | 110 | gp15 | 93 | 4.84 | 52 | 2.26 |
| HTJP | gp50 | 100 | gp16 | 89 | 6.99 | 49 | 1.57 |

| **80α/SaPI1** | | **vs.** | **Lambda** | **PDB ID: 8K38** | | | |
|---|---|---|---|---|---|---|---|
| | Gene product | # res | gp | total | RMSD | pruned | RMSD |
| HTCP | gp49 | 110 | gpW | 49 | 6.59 | 23 | 2.34 |

| **80α/SaPI1** | | **vs.** | **Lambda** | **PDB ID: 8K37** | | | |
|---|---|---|---|---|---|---|---|
| Protein | Gene product | # res | gp | total | RMSD | pruned | RMSD |
| HTJP | gp50 | 100 | gpFII | 84 | 14.43 | 25 | 2.28 |
| TrP | gp52 | 127 | gpU | 114 | 5.42 | 48 | 2.34 |

| **80α/SaPI1** | | **vs.** | **HK97** | **PDB ID: 3JVO** | | | |
|---|---|---|---|---|---|---|---|
| Protein | Gene product | # res | gp | total | RMSD | pruned | RMSD |
| HTCP | gp49 | 110 | gp6 | 85 | 3.61 | 64 | 1.66 |

| **80α/SaPI1** | | **vs.** | **JBD30** | **PDB ID: 8RKB** | | | |
|---|---|---|---|---|---|---|---|
| Protein | Gene product | # res | gp | total | RMSD | pruned | RMSD |
| HTCP | gp49 | 110 | gp41 | 81 | 5.21 | 55 | 1.78 |

| **80α/SaPI1** | | **vs.** | **GTA** | **PDB ID: 6TE9** | | | |
|---|---|---|---|---|---|---|---|
| Protein | Gene product | # res | gp | total | RMSD | pruned | RMSD |
| HTCP | gp49 | 110 | Rcc01688 | 58 | 23.31 | 28 | 1.05 |
| HTJP | gp50 | 100 | Rcc01689 | 91 | 7.25 | 53 | 1.61 |
| TrP | gp52 | 127 | Rcc01690 | 110 | 10.86 | 49 | 1.62 |

¶Total=total number of residue pairs compared. The RMSD value to the right is between this number of residue pairs.

§Pruned=number of residue pairs compared after pruning to remove all pairs with an RMSD>3.5Å, and the corresponding RMSD value. Aligned in UCSF Chimera using the BLOSUM-62 matrix and 50% weight on secondary structure vs. sequence.
